# Thalamic activity during scalp slow waves in humans

**DOI:** 10.1101/2021.10.11.463988

**Authors:** Péter P. Ujma, Orsolya Szalárdy, Dániel Fabó, Loránd Erőss, Róbert Bódizs

## Abstract

Slow waves are major pacemakers of NREM sleep oscillations. While slow waves themselves are mainly generated by cortical neurons, it is not clear what role thalamic activity plays in the generation of some oscillations grouped by slow waves, and to what extent thalamic activity during slow waves is itself driven by corticothalamic inputs. To address this question, we simultaneously recorded both scalp EEG and local field potentials from six thalamic nuclei (bilateral anterior, mediodorsal and ventral anterior) in fifteen epileptic patients (age-range: 17-64 years, 7 females) undergoing Deep Brain Stimulation Protocol and assessed the temporal evolution of thalamic activity relative to scalp slow waves using time-frequency analysis. We found that thalamic activity in all six nuclei during scalp slow waves is highly similar to what is observed on the scalp itself. Slow wave downstates are characterized by delta, theta and alpha activity and followed by beta, high sigma and low sigma activity during subsequent upstates. Gamma activity in the thalamus is not significantly grouped by slow waves. Theta and alpha activity appeared first on the scalp, but sigma activity appeared first in the thalamus. These effects were largely independent from the scalp region in which SWs were detected and the precise identity of thalamic nuclei. Our results indicate that while small thalamocortical neuron assemblies may initiate cortical oscillations, especially in the sleep spindle range, the large-scale neuronal activity in the thalamus which is detected by field potentials is principally driven by global cortical activity, and thus it is highly similar to what is observed on the scalp.

## Introduction

Due to the difficulty of accessing subcortical structures, the electroencephalogram (EEG) in sleeping humans is most frequently recorded from the scalp, and even invasive recordings are often limited to subdural grids. The thalamus is an important pacemaker and generator of sleep EEG oscillations (David et al. 2013; Fuentealba and Steriade 2005; Steriade 2003), but human thalamic recordings in sleep are rare.

Several lines of evidence point at the significance of thalamic activity in orchestrating sleep EEG oscillations. First, while slow waves exist in deafferented cortical tissue (Timofeev et al. 2000), thalamic activity plays a crucial role in regulating their rhythm (Crunelli and Hughes 2010; David et al. 2013). Second, sleep spindles arise in the entirety of the thalamocortical system, with a crucial role of the reticular thalamus, which is hypothesized to mainly regulate the activity of thalamocortical cells (Fernandez and Lüthi 2020; Fuentealba and Steriade 2005; Steriade 2003). Third, slow waves synchronize spindles and spindles synchronize hippocampal ripples (Clemens et al. 2007; Clemens et al. 2011; Staresina et al. 2015), and thalamic activity during spindles is hypothesized to regulate hippocampal input to the cortex in the process of consolidating recent memory traces (Peyrache et al. 2011). While oscillations beside slow waves, spindles and ripples are less often studied, some findings suggest a role of thalamic regulation in the theta (Gonzalez et al. 2018) and alpha (Halgren et al. 2019) oscillations as well.

While human thalamic recordings are not routinely performed, some previous research is available about human thalamic activity during sleep. A systematic study of invasive recordings from the human thalamus was reported already in the 1940s (Williams and Parsons-Smith 1949). Some further studies of sleep oscillations (Velasco et al. 2002; Velasco et al. 1979; Wennberg et al. 2002) and patterns of single-cell activity (Tsoukatos et al. 1997) in this structure were published decades ago. More recently, a study of three epileptic patients (Tsai et al. 2010) reported significant coherence at delta, theta and sigma frequencies between the cortex and the thalamus, suggesting the synchronous involvement of both structures in the generation of these oscillations. A study of pulvinar recordings (Mak-McCully et al. 2017) showed that in the thalamus downstates follow, but spindles precede analogous cortical activity, suggesting the thalamic origin of spindles, but the cortical origin of slow-wave activity (Csercsa et al. 2010; Nir et al. 2011; Tononi and Cirelli 2014). In the same patients, another study (Gonzalez et al. 2018) investigated events surrounding downstates, and reported an increase of low-frequency activity (including oscillatory theta) during and high-frequency activity after the downstate. A further study (Bastuji et al. 2020) studied sleep spindle activity in a wide variety of thalamic nuclei and found that – beside frequent co-occurrence with thalamic spindles – local spindles are also present in the thalamus. In this study, the precise temporal pattern of cortical and thalamic spindle occurrence was not studied.

In our study, we aimed to systematically describe thalamic field potentials during slow waves reported on the cortex. For this, we used a large patient population (N=15), recordings from a total of six different precisely localized nuclei with a bipolar reference to focus on local activity (Wennberg et al. 2002), and time-frequency analysis at a wide array of frequencies. We find that even with bipolar reference, thalamic activity at the field potential level is surprisingly similar to cortical activity, with only small time lags and little dependence on the localization of cortical slow waves. However, we did find that slow (delta-alpha) and beta activity slightly but significantly lags cortical activity in the thalamus, but the opposite pattern is seen for sigma activity reflecting sleep spindles.

## Methods

### Patients

15 pharmacoresistant, surgically non-treatable epilepsy patients (M_age_=36.9 years, range: 17-64; 7 females) participated in an anterior thalamus deep brain stimulation (DBS) protocol at the National Institute of Clinical Neurosciences, Budapest, Hungary. The research was approved by the ethical committee of the National Institute of Clinical Neurosciences. Patients signed informed consent for participating in the study. Detailed clinical and demographic data of this patient population are reported elsewhere (Szalárdy et al. 2021).

### Implantation procedure and clinical recording protocol

During general anesthesia, a pair of quadripolar Medtronic DBS electrodes (contact lengths 1.5 mm, intercontact spacing 0.5 mm, electrode diameter 1.27 mm) was bilaterally and stereotaxically inserted into the thalamus, targeting the anterior nucleus. Depending on patient characteristics and according to the decision of the neurosurgical team, a frontal transventricular, extraventricular or posterior parietal extraventricular approach was used for electrode placement. After the surgery, patients underwent a 48 hour video-EEG monitoring protocol with co-registered thalamic LFP (externalized thalamic leads) and scalp EEG/polysomnography, with no thalamic stimulation in the first 24 hours, yielding a full night of stimulation-free recordings which were used for analysis.

Thalamic contacts were localized with high precision using preoperative MRI and postoperative CT images co-registered with the FMRIB Software Library (FSL, Oxford, FLIRT), and double-checking by using the mamillothalamic tract as an anatomical guide. (See also (Simor et al. 2021) for co-registration details). With this procedure, we identified thalamic contacts recording from three different thalamic nuclei or regions, the anterior thalamus (ANT), the mediodorsal nucleus (MD) or the ventral anterior nucleus (VA) on either the left or the right side for a total of six possible thalamic nuclei/regions (Table 1). For simplicity, we refer to all of these regions as ‘nuclei’, although the ANT is technically composed of several subnuclei.

**Table 1.**
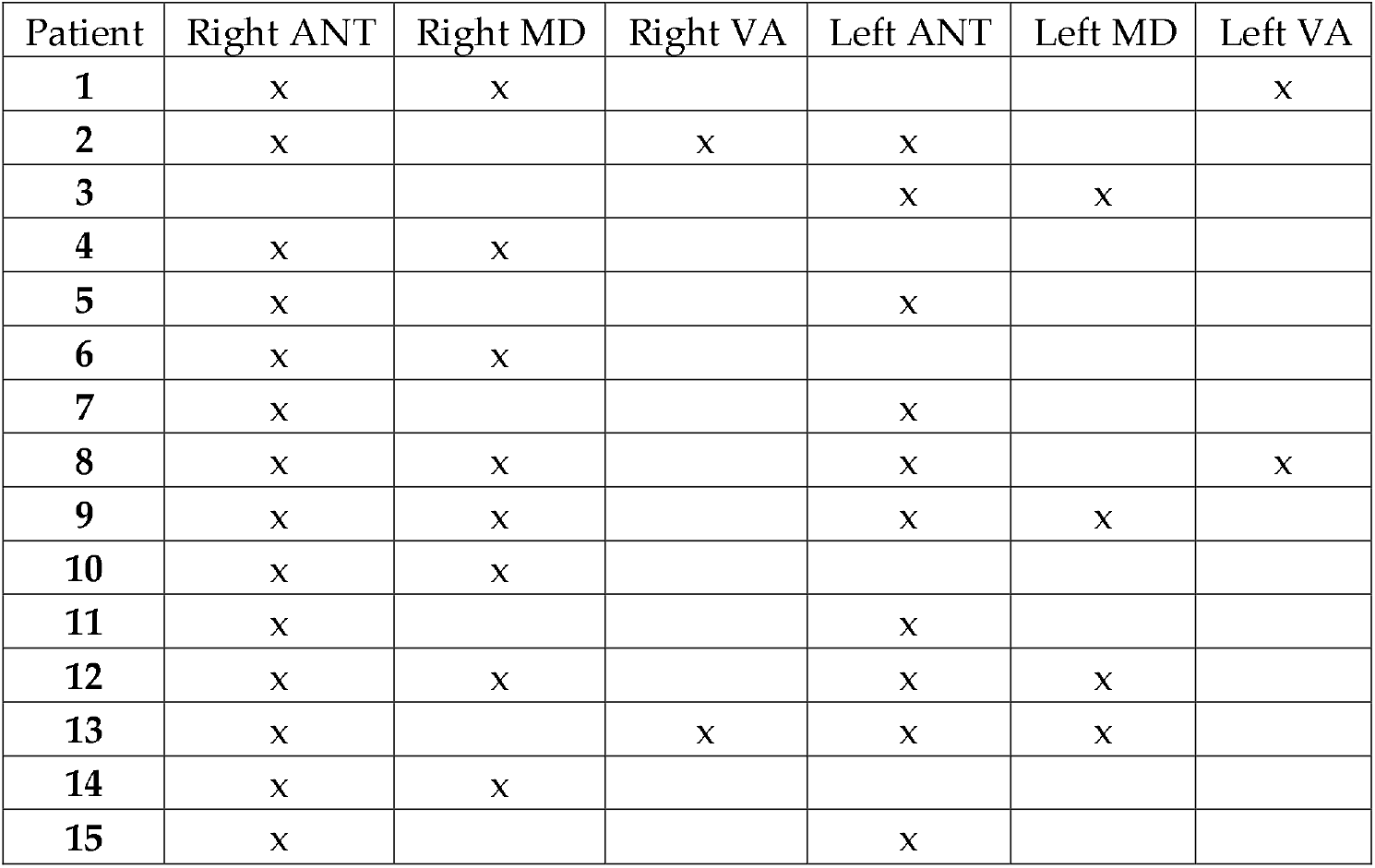
Availability of data from each patient and thalamic nucleus.

### Electrophysiology

Scalp EEG and thalamic LFP signals were recorded by an SD-LTM 64 Express EEG/polygraphic recording system. Physiological signals were recorded at 8192 Hz/channel effective sampling rate with 22 bit precision and hardware input filters set at 0.02 (high pass: 40 dB/decade) and 450 Hz (low pass: 40 dB/decade), firmware-decimated by a factor of 4 for a final sampling rate of 2048 Hz.

Scalp EEG was recorded according to the international 10-20 system, extended with the inferior temporal chain and two anterior zygomatic electrodes, which, however, we did not use for these analyses. We considered the following 16 common electrodes in each patient: Fp1 Fp2 F3 F4 F7 F8 C3 C4 P3 P4 O1 O2 T3 T4 T5 T6. Missing (5 channels in 1 patient) or artefactual (10 channels in 6 patients) scalp channels were treated as missing data. All electrodes were referenced to CP1, with the ground placed at CP2. Submental electromyograms (EMGs) were recorded by bipolarly referenced electrodes placed on the chin. Scalp electrodes were re-referenced to the mean of two channels from the lower temporal chain (Tp9 and Tp10, the latter replaced by the adjacent Po10 for Patient 7 due to persistent noise on Tp10) before analysis, approximating a mastoid reference.

Thalamic LFP signals were recorded with a bipolar reference between adjacent contacts. We considered data to be available from a thalamic nucleus if there were two contacts localized inside of it or if one contact was inside of it and another in adjacent tissue (Deutschová et al. 2021).

All-night sleep records were scored according to standard AASM criteria on a 20 seconds basis (Berry et al. 2015) by an expert. Artifact-contaminated data segments were visually marked on a 4 seconds basis and excluded from analysis.

### Slow wave detection

Slow waves (SWs) were automatically detected on each scalp channel based on previously established criteria (Menicucci et al. 2009). After pre-filtering NREM sleep EEG data to 0.5-4 Hz, slow waves were defined as negative deflections of a duration between 200 and 1000 msec and a minimum amplitude of - 37.5 μV and a minimum peak-to-peak amplitude of 75 μV relative to the subsequent positive half-wave. An alternative detection criterion with minimum negative deflections at −75 μV and a minimum peak-to-peak amplitude of 140 μV, while resulting in slightly more valid detections in a healthy sample (Ujma et al. 2019), was investigated but rejected because it resulted in zero detections in several patients and channels.

For our analyses, we needed to precisely measure the temporal offset between scalp SWs and thalamic activity. Therefore, we restricted analyses for scalp channels where at least 20 SWs were detected so that thalamic activity could be averaged based on a sufficient number of events. We also excluded data from individual combinations of scalp channels, thalamic nuclei, subjects and frequency bands based on visual inspection of the resulting average thalamic waveform. This resulted in a total of 37 rejections (out of a total of 596 patient-nucleus-channel combinations per frequency band) for the delta band, 65 for the theta, 91 for the alpha, 58 for the low sigma, 77 for the high sigma, 89 for the beta and 1 for the gamma range. Because the thalamic waveform in the gamma range tended to be highly irregular and not significantly coupled to SWs (see Results), we did not attempt to improve data quality by more exclusions from this frequency range.

For some analyses, we report data from scalp regions instead of individual channels. In these cases, results from all channels within the same region were averaged. The regions and constituent channels were: frontal (Fp1, Fp2, F3, F4), centro-parietal (C3, C4, P3, P4), right temporal (F8, T4, T6), left temporal (F7, T3, T5) and occipital (O1, O2).

We initially detected and analyzed SWs separately for each scalp channel. However, we also described SWs as propagating events, which allowed the analysis of thalamic activity relative to the specific channel SWs originated from. For this, we used a method based on previous literature (Menicucci et al. 2009; Ujma et al. 2018). We first detected SWs on individual channels. Next, we ordered SWs by the timing of maximal negative deflections. The first SW was assumed to be a canonical wave (Menicucci et al. 2009), initiating a propagating event from the channel it was detected on and all other SWs occurring within 200 msec were assumed to belong to this event, with corresponding propagation lags on their respective channels of origin. The first SW outside this 200 msec window was assumed to initiate the next propagating event consisting of all other SWs within a new 200 msec window. This process was repeated until all SWs were grouped. The characteristics of propagating SWs obtained with this method were highly similar to what was reported in previous literature (see Results).

### Thalamic data analysis

In each patient, the signal of all available thalamic channels was triggered to the peak of the negative deflection of each slow wave on each available scalp channel (time window: [−2000 2000] msec relative to the scalp SW peak). The time-frequency decomposition of these thalamic signals was calculated using the cwt() function of the MATLAB Signal Processing Toolbox. The resulting instantaneous amplitudes were log-transformed and only the [−1000 1000] msec section was retained to eliminate edge artifacts. In order to correct for baseline effects in the thalamogram, the thalamic signal around a random time point selected 2-3 seconds before or after each SW peak was subjected to the same time-frequency decomposition and the final thalamic time-frequency signal around each scalp SW was calculated as the difference between the instantaneous amplitudes of the actual and the adjacent random signal. Instantaneous amplitudes were averaged across thalamic channels within the same patient and thalamic nucleus to yield a single within-nucleus measure. A sample of 10000 randomly selected data points in artifact-free NREM sleep was also selected, and the thalamic time-frequency signal around these points was calculated as described above in each available nucleus. To establish a baseline thalamic activity, we randomly selected as many of these random time-frequency signals as there were actual SWs on each scalp channel in each patient. In statistical analyses, we compared instantaneous amplitudes of the thalamic signal during scalp SWs to this baseline. Statistical significance was determined using two-sample t-tests. This analysis was performed individually for each patient, scalp channel, thalamic nucleus, time-frequency time point and frequency, the comparison baseline always being the thalamic instantaneous amplitude in the same patient, in the same thalamic nucleus and at the same time-frequency time point and at the same frequency during an identical number of randomly selected NREM time points. We averaged results across patients, but instead of estimating a meta-analytic statistical significance of this average effect size we more conservatively converted p-values to z-values, averaged them across patients and transformed them back. This method is similar to Fisher’s method of averaging the natural logarithm of p-values (Mosteller and Fisher 1948). It can be interpreted as a meta-analysis with the null hypothesis that thalamic activity triggered to SWs is not different from random data in individual patients, while the null hypothesis of an ordinary meta-analysis would be that the average activity across all patients is not significantly different from random data. We used the same method of averaging both effect sizes and p-values when averaging results across scalp channels or frequencies to reduce the complexity and redundancy of the results for illustration. Figure 1 illustrates this process.

**Figure 1.**
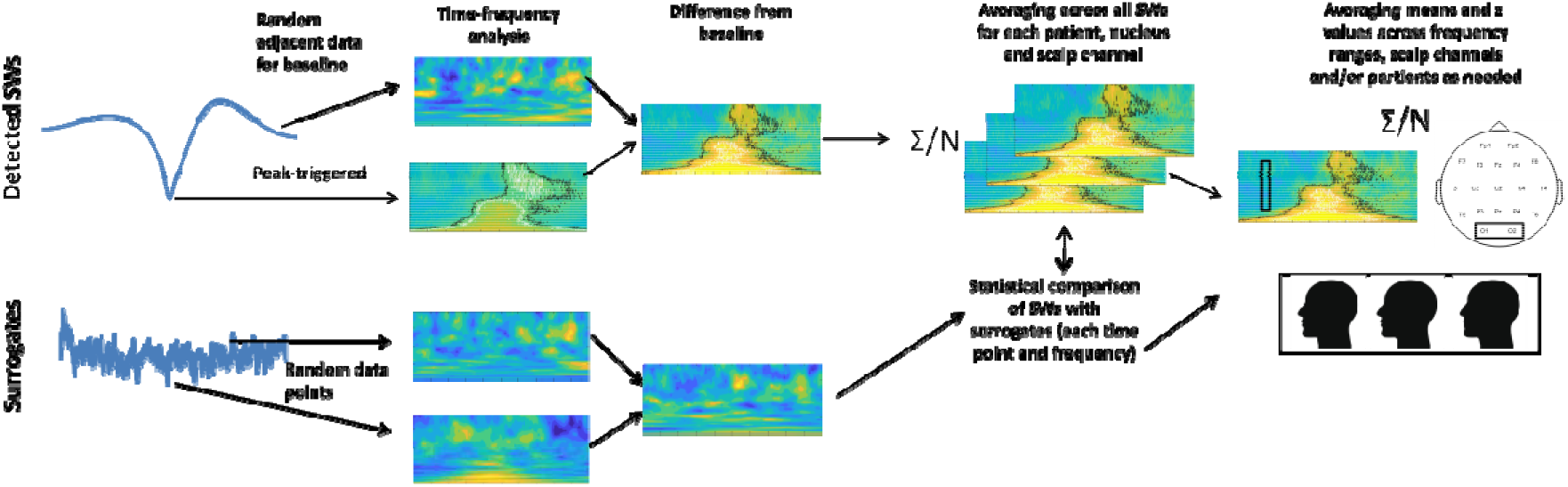
Illustration of the data analysis pipeline.

We estimated the lag between scalp SWs and thalamic activity between different nuclei and scalp channels by averaging the time course of thalamic activity within each frequency band across all SWs finding the relative timing of the largest deviation from zero. This was estimated for each patient, each frequency band, each available scalp channel and each thalamic nucleus. For the sigma frequency band, we restricted the search for largest deviations to time points after the scalp SW peaks to avoid the spurious detection of peaks resulting from the SW waveform itself (see Results). Lag relative to the scalp SW peak was modeled with a mixed linear model (implemented in the MATLAB lmer() function) as a function of frequency band, scalp channel and thalamic nucleus with random intercepts by patient. We explored changes in model fit after adding interactions (band*scalp region, band*nucleus, nucleus*scalp region) and comparing model fit using the −2 Log-likelihood (−2LL) and Akaike Information Criterion AIC (AIC) statistics. −2LL is the maximum likelihood analog to residual sum of squares in ordinary least squares models, and it indicates deviance (variance unexplained by the model). AIC is derived from −2LL by introducing a penalty term for more complex models. If an independent variable has minimal utility, its inclusion may improve −2LL but lead to increased AIC values, indicating that the explanatory power of this variable is offset by the increased model complexity caused by its inclusion. A significantly better model fit based on both −2LL and AIC was interpreted as evidence for the presence of an interaction.

## Results

The thalamic time-frequency around scalp SWs was similar to previous descriptions of other thalamic nuclei (Gonzalez et al. 2018) or the scalp EEG itself (Muehlroth and Werkle-Bergner 2020). An increase in low delta-frequency activity was observed around scalp SWs even in the thalamus, with a more defined increase in theta and alpha-frequency activity immediately preceding and during the peak. 100-600 msec after the scalp SW peak, an increase of broadly defined beta and sigma-frequency activity (10-20 Hz) was seen with a progressively decreasing frequency, corresponding to beta activity and sleep spindles during the slow wave upstate. Figure 2 illustrates this pattern using average data in the left ANT nucleus during SWs detected on Fp1, but very similar results were observed in all scalp channel-thalamic nucleus pairing, and individually in each patient (detailed results individually for all patients, nuclei and scalp channels are available in the Supplementary data).

**Figure 2.**
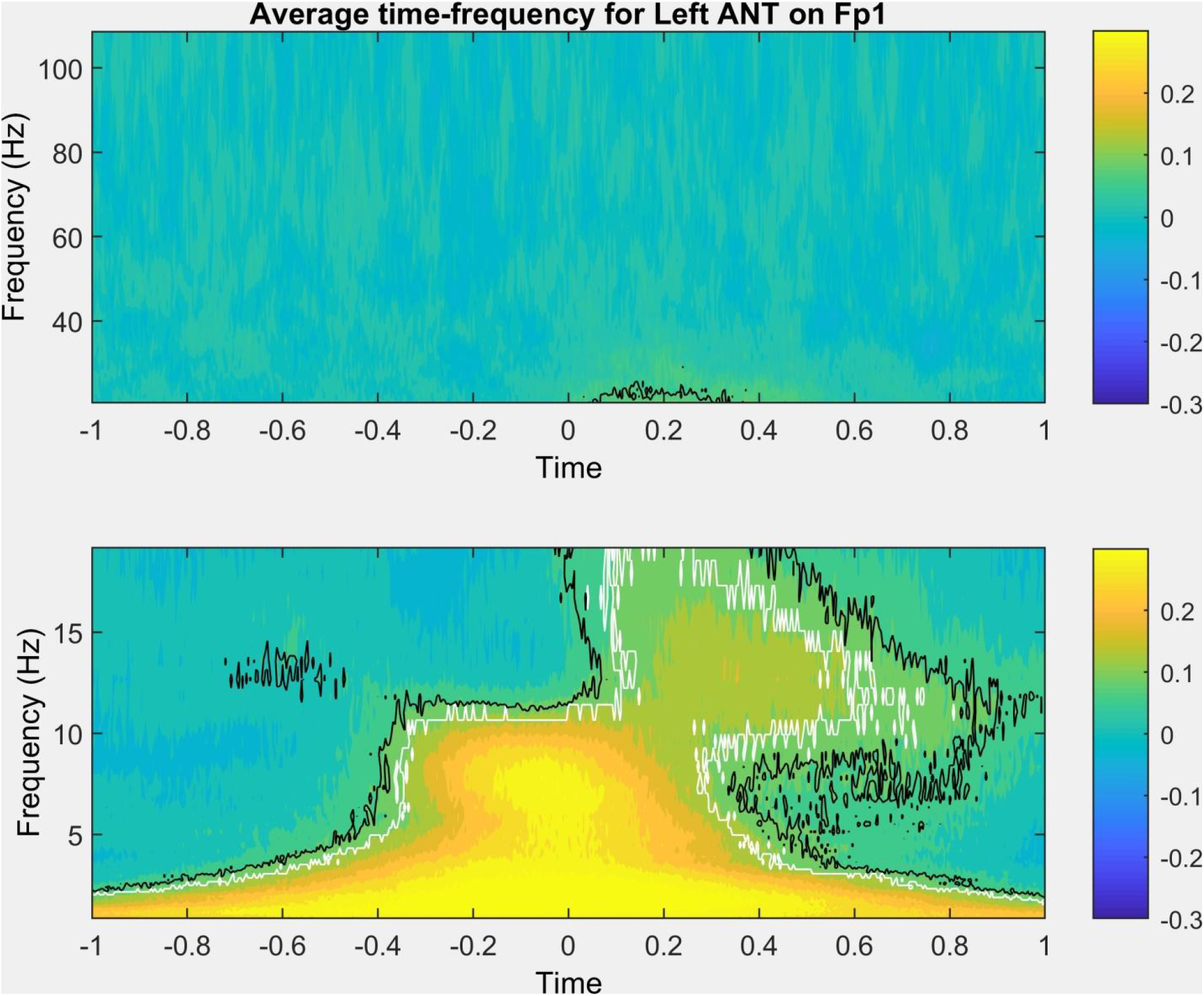
Representative time-frequency plot (data averaged across all patients, representing average field potentials in the left anterior thalamus, triggered to the negative peak of slow waves detected on Fp1). Black lines trace time and frequency values where activity was significantly more extreme than in random segments at uncorrected p<0.05. White lines trace time and frequency values where the effects also survived correction for the false discovery rate. The bottom and top subplots show <20 Hz and >20 Hz activity, respectively, with no data gap but a break in scale for optimal visibility.

Aggregate results by thalamic nucleus (including the scalp for illustration, based on time-frequency analysis of the scalp EEG data of the same channel the SW was detected on), frequency band and region are shown on Figure 3. Note the similar shape of the curves across nuclei and their similarity to the scalp, but notable differences between frequency bands.

**Figure 3.**
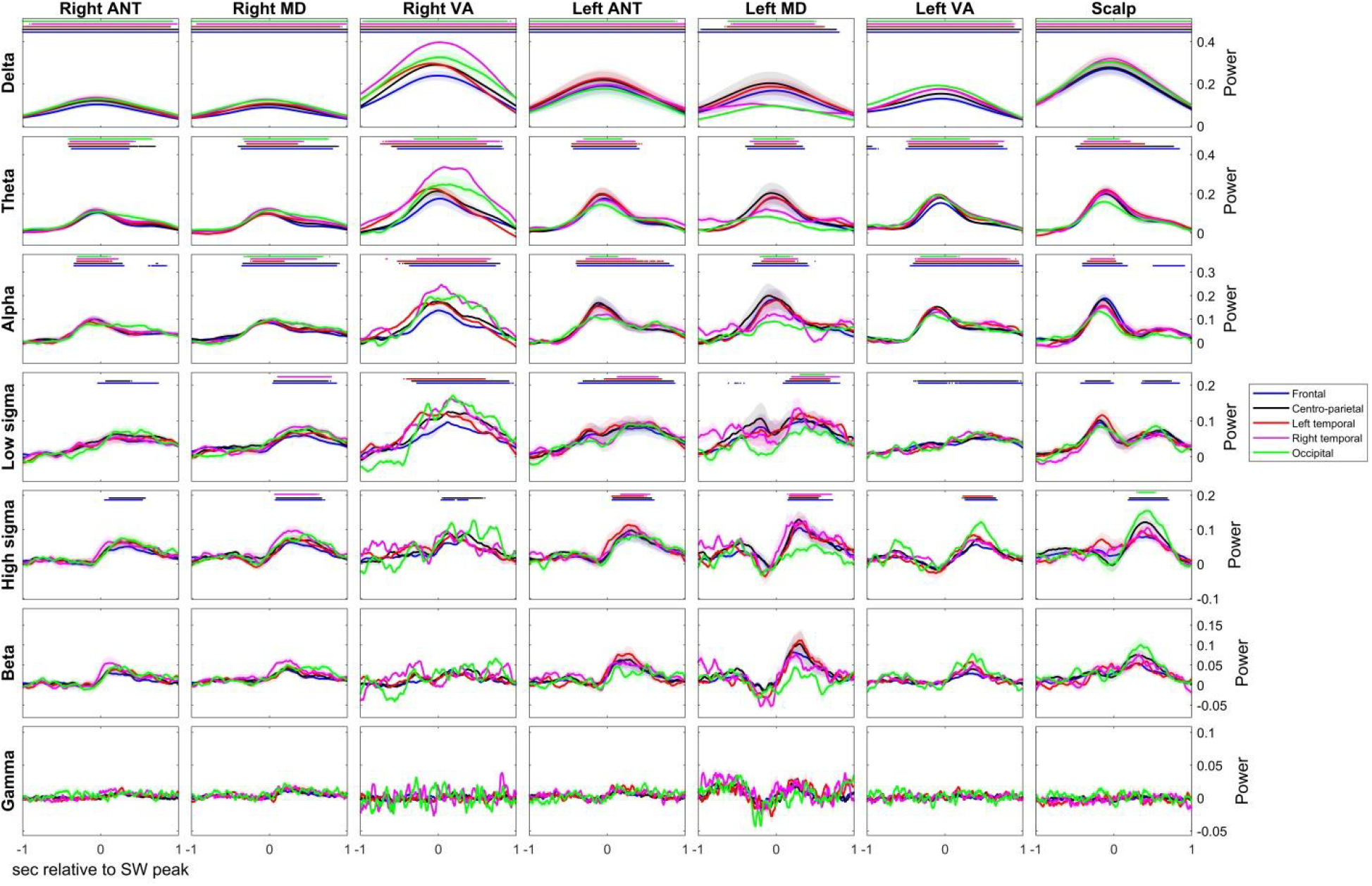
Average time-frequency activity triggered to scalp SW peaks in six thalamic nuclei by frequency band and scalp region. All SWs detected on a channel in the target region were considered for this analysis. The course of time-frequency activity on the scalp itself is shown for reference. Dots indicate that time-frequency activity is significantly different (after FDR correction) from the baseline established in random samples. Because a different subset of patients provided data for each of the six nuclei, between-nucleus power differences also reflect between-patient differences in signal voltage and cannot be directly compared. Shades around the lines indicate 95% confidence intervals across patients. Note that 95% confidence intervals do not imply statistical significance (which is established by averaging log p-values obtained in each patient by comparing actual activity to random samples) but how divergent the course of time-frequency activity is across patients.

Delta, theta and alpha-frequency activity in the thalamus coincided with or slightly preceded the negative peak of the scalp SW. Conversely, in ~200-300 msec the negative peak was followed by an up-state characterized by beta, then high sigma, then finally low sigma-frequency oscillations. The maximum of the thalamic gamma band did not exhibit a characteristic temporal relationship to the scalp SW peak. No clear topographic pattern was seen: thalamic activity related to scalp SWs in a similar way regardless of which region they were detected in. The mean lag of the envelope peak in each nucleus and at each frequency band relative to the scalp SW peaks is shown on Figure 4. Comparative data from the scalp - that is, from the detected SW itself - is also shown.

**Figure 4.**
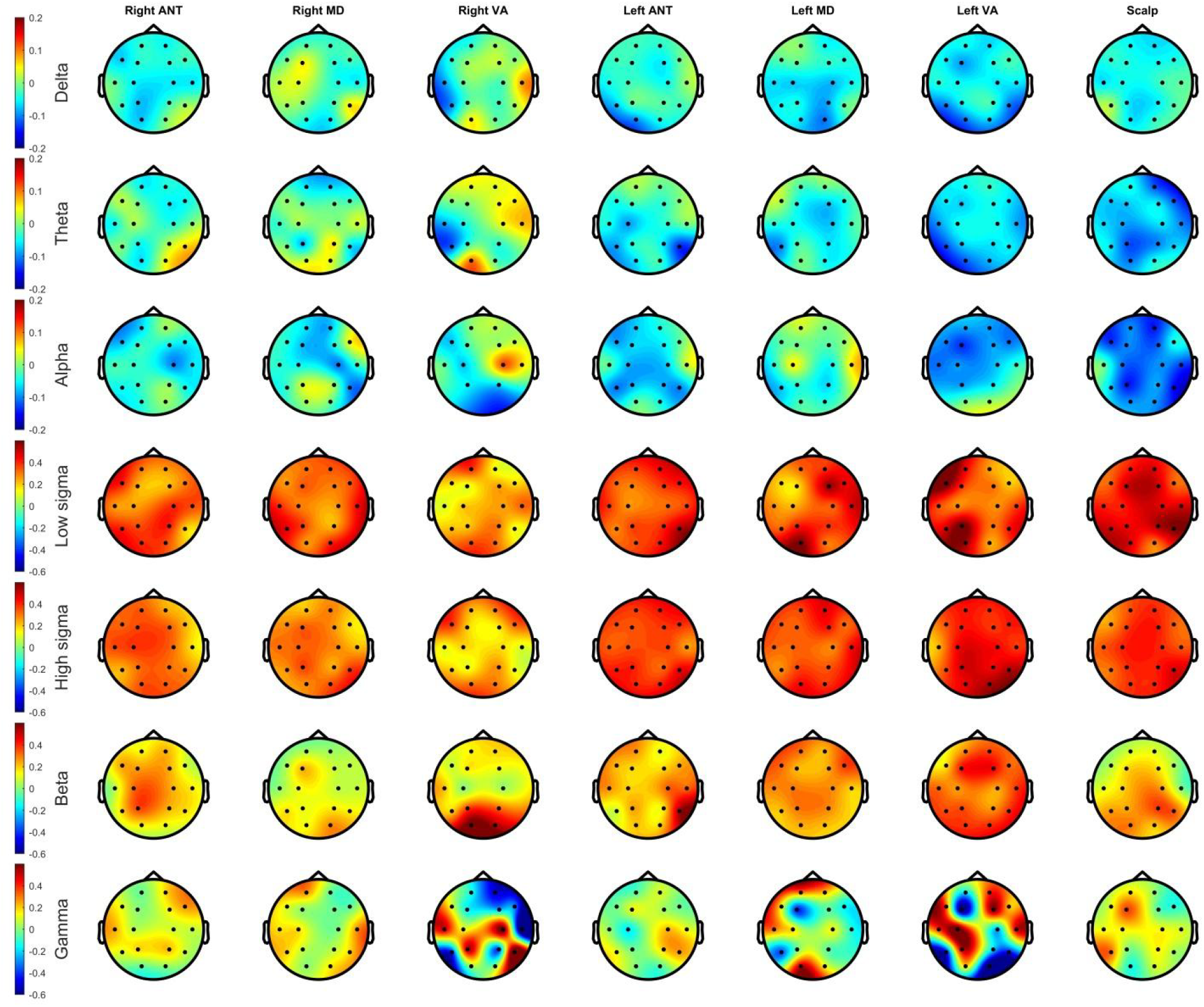
Temporal lag (in seconds) of activity in each frequency band relative to SW negative peaks as a function of thalamic nucleus and scalp region. Data from the scalp itself is shown for reference.

The analyses previous analyses established the temporal patterns of various frequencies relative to the negative scalp SW peak. Note that this is not identical to activity on the scalp, and a negative lag in the thalamus does not automatically imply that thalamic activity occurred earlier than scalp activity. For example, the peak of delta power is observed slightly before the negative SW peak not just in the thalamus, but even on the scalp itself (Figure 4). In a more precise analysis we aimed to directly compare temporal lags from various data sources. For this, we used a linear mixed model (implemented with the MATLAB lmer() function, with an identity link function and assumed normal distribution) to model time lags of maximal thalamic activity relative to scalp SW peaks (effectively the data shown on Figure 4) as a function of thalamic nucleus, scalp region of SW origin and frequency band, with random intercepts by patient. The model intercept (the estimated mean lag of the reference category, delta activity in the scalp during frontal SWs) was −0.04 seconds (p=0.02). Of the other effects, only band effects were significant: gamma (0.11 sec, p=10^−19^), beta (0.23 sec, p=10^−64^), high sigma (0.38 sec, p=10^−162^) and low sigma (0.41 sec, p=10^−187^) activity followed SW peaks with significant lags. Adding band*nucleus interactions – assuming that the lag estimated in each nucleus is affected by the frequency band being investigated – significantly improved model fit by both the −2LL and AIC measures (Likelihood ratio=106.12, p=10^−9^; ΔAIC=−34.13). Also adding region*nucleus interactions – assuming that SWs originating in different scalp areas result in different lags in the thalamic nuclei – did not improve model fit any further (Likelihood ratio=22.23, ΔAIC=25.78). Adding region*band interactions to the model also containing nucleus*band interactions – assuming that SWs originating in different scalp areas result in different lags in different frequency bands – improved model fit, but resulted in a lower AIC value due to loss of parsimony (Likelihood ratio= 37.628, p=0.037; ΔAIC=10.38). Therefore, our preferred model remained the one with main effects and band*nucleus interactions. Consequently, we also ran our models by frequency band, but not by scalp region or nucleus. All model coefficients are illustrated on Figure 5.

**Figure 5.**
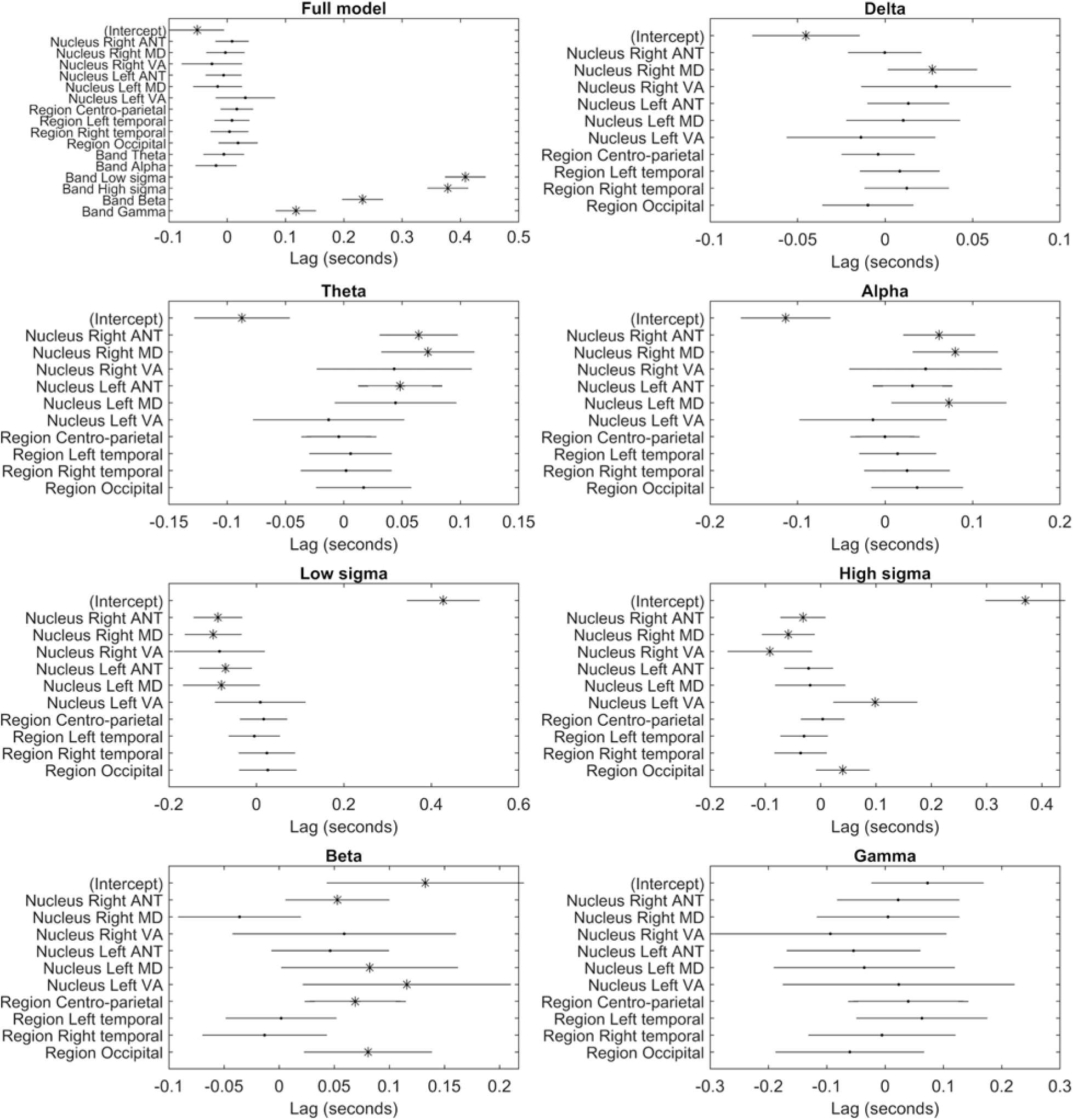
Estimated mean lag of maximum activity relative to scalp SW peaks in the baseline model using all data (first panel) and separately by frequency band (other panels). The intercept in each model can be interpreted as the estimated mean of the reference category (frontal delta activity on the scalp in the baseline model, frontal activity on the scalp in the other models). Model coefficients for the other categories can be interpreted as the estimated difference of the mean lag of that category from the intercept. Asterisks indicate significant coefficients (an intercept significantly different from zero or a category in which lag is significantly different from the reference category indicated by the intercept). Error bars indicate 95% confidence intervals.

The most important result of these models is that in the theta and alpha frequency ranges, SW peak-triggered scalp activity (indicated by the intercept) preceded thalamic activity by about 50-100 msec. This effect was similar in all thalamic nuclei except the left VA, but it was imprecisely estimated for VA nuclei where data was only available from 2-2 patients from both sides. A smaller scalp advantage remained a trend in the delta band, significant only in the right MD nucleus.

The inverse was seen in the low and high sigma frequency range: here, thalamic activity preceded scalp activity by 50-100 msec, again with the exception of the left VA. The difference was largest and significant for all nuclei (except VA) for the low sigma range, while in the high sigma range it was smaller and only significant for the right ANT, MD and VA, remaining a trend for the left ANT and MD and a significant opposite pattern for the left VA.

In the beta range, thalamic activity again generally followed scalp activity by ~50-150 msec. This remained a trend in the right VA and the left ANT and an opposite trend was seen in the right MD.

A weakness of our analyses so far has been that we pooled all SWs detected on each scalp channel. SWs are known to be propagating events (Massimini et al. 2004; Menicucci et al. 2009; Ujma et al. 2018), and therefore this method mixes SWs genuinely originating from cortical tissue underlying scalp channels with SWs merely propagating there, which may bias temporal lags observed in the thalamus if corticothalamic propagation occurs from the first cortical area exhibiting the neural firing patterns underlying SWs. Therefore, we repeated our analyses using only the original SWs of each scalp channel – SWs which have been demonstrated to appear on that channel first in a chain of propagating events (see Methods). A disadvantage of this strategy is that it drastically reduces the number of available SWs on each channel, because each SW can appear on multiple channels in the initial analysis but by definition can originate from a single one in this new one. Figure 6 illustrates the features of propagating SWs. As reported in previous literature (Massimini et al. 2004; Menicucci et al. 2009; Ujma et al. 2018), the vast majority of SWs originated in frontal channels and these also had the largest amplitudes, SWs were most likely to propagate to adjacent channels and SWs originating in occipital channels were the most likely to propagate forward while frontal SWs often remained local.

**Figure 6.**
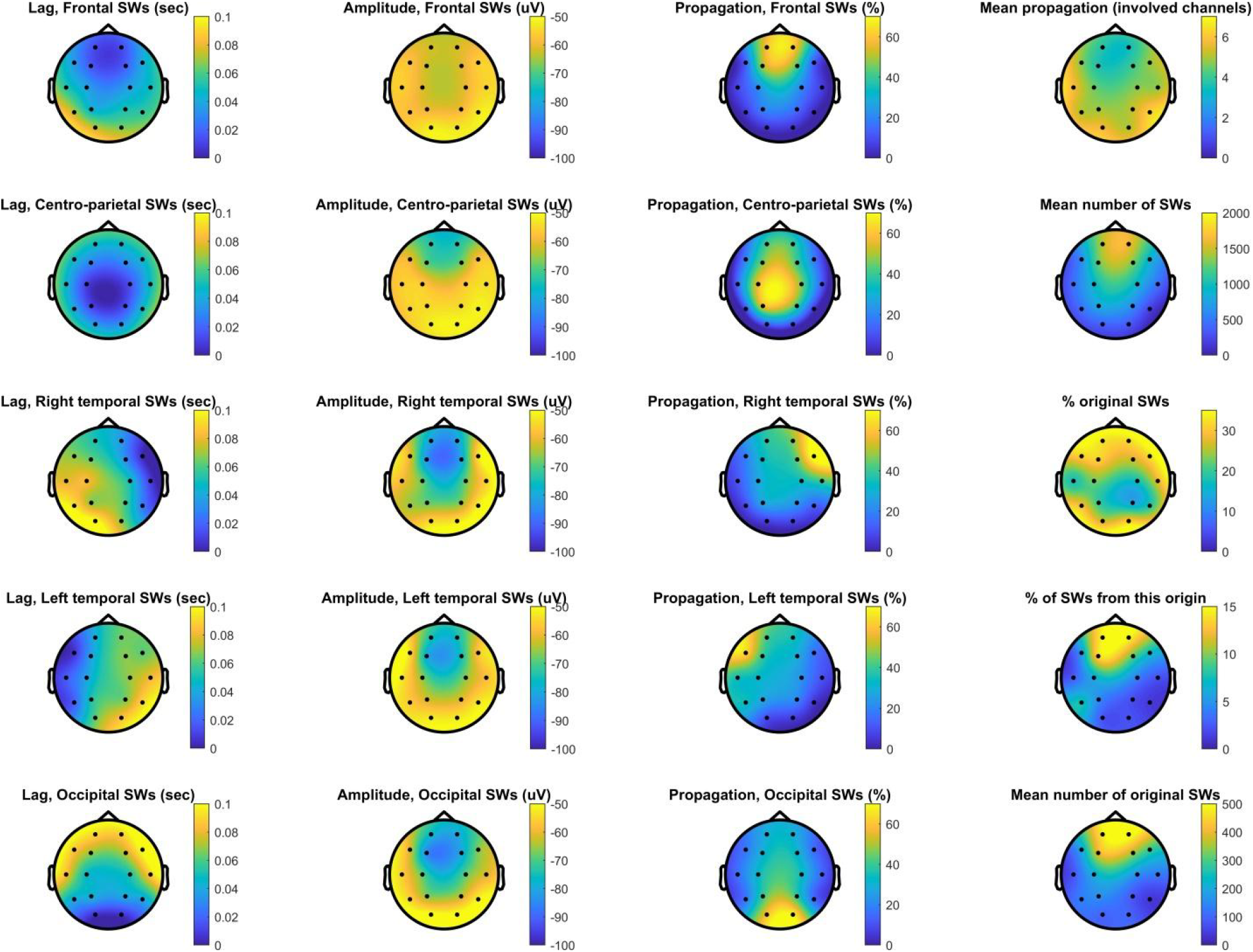
Characteristics of SW propagation. This plot shows the topographical distribution of propagation probability as a function of region of origin (first column), the mean amplitude of propagated SWs (second column), and the percentage of SWs detected in the region of origin followed by a propagated SW on other channels (third column). In the fourth column, we show the topographical distribution of the mean number of other channels SWs detected on each channel propagated to (top plot), the mean number of SWs detected on each channel (second from top), the percentage of original SWs relative to all SWs detected (third from top), the percentage of all SWs originating from each channel (fourth from top) and the mean number of original SWs (bottom plot). All data is averaged across patients.

Similar results were seen when only original waves were used (Figure 7). Scalp SWs downstates were accompanied by delta, theta and alpha frequency activity in all thalamic nuclei, without obvious regional or nucleus effects. Beta and sigma activity followed SW peaks by 200-300 msec, while there was no clear pattern for gamma activity. Due to our stringent definition of statistical significance, however, these effects were only significant in the lower frequency ranges and relative to frontal (and in one case, centro-parietal) SWs.

**Figure 7.**
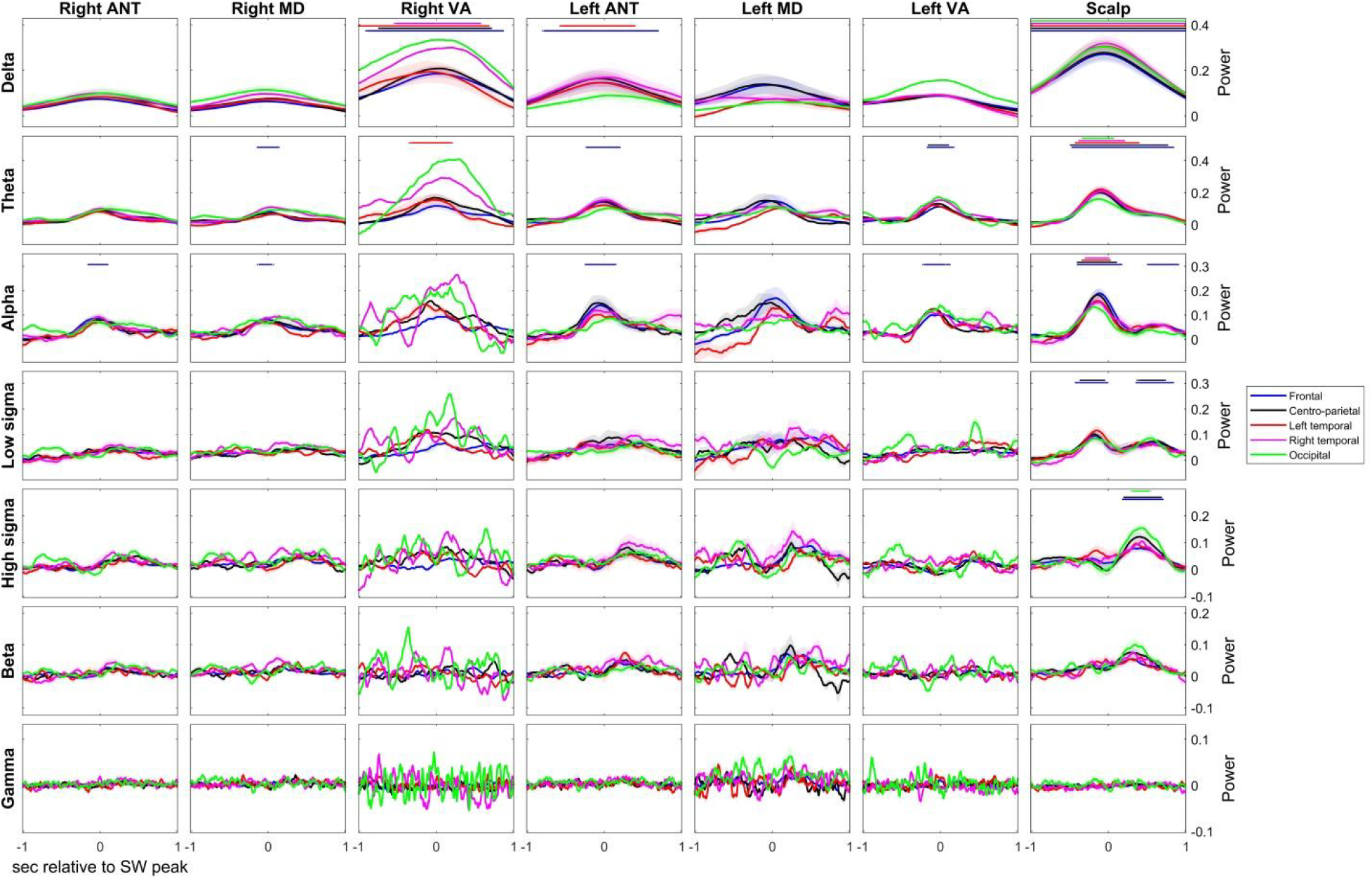
Average time-frequency activity triggered to original scalp SW peaks in six thalamic nuclei by frequency band and scalp region. Only SWs originating from a channel in the target region were considered for this analyses. The course of time-frequency activity on the scalp itself is shown for reference. Dots indicate that time-frequency activity is significantly different (after FDR correction) from the baseline established in random samples. Because a different subset of patients provided data for each of the six nuclei, between-nucleus power differences also reflect between-patient differences in signal voltage and cannot be directly compared. Shades around the lines indicate 95% confidence intervals across patients. Note that 95% confidence intervals do not imply statistical significance (which is established by averaging log p-values obtained in each patient by comparing actual activity to random samples) but how divergent the course of time-frequency activity is across patients.

Because of the low number of available SWs on most non-frontal channels, lag could not be estimated with sufficient precision to be statistically modeled (Supplementary figure S1).

## Discussion

Our study is the first to systematically measure thalamic field potentials during scalp SWs. Our investigation involved six thalamic nuclei and pooled data from a total of fifteen patients. We could demonstrate that time-frequency activity of thalamic field potentials is generally similar to what is observed on the scalp. All thalamic nuclei exhibit clear downstates when an analogous phenomenon occurs on the scalp, characterized by low-(delta-alpha) frequency activity. This is followed by a clear upstate in ~150-600 msec (Figure 1), characterized by sequential beta, high sigma and low sigma activity, in a manner very similar to what is observed on the scalp (Muehlroth and Werkle-Bergner 2020) and in the pulvinar (Gonzalez et al. 2018). Only slight differences relative to the scalp were seen in the timing of these phenomena. Low-frequency (especially in the theta and alpha ranges) activity and beta activity in the thalamus slightly trails scalp activity. An opposite pattern was seen in the sigma range, where – especially in the low sigma range – thalamic activity preceded scalp activity. We observed no systematic difference between thalamic nuclei in these trends. Surprisingly, we also did not observe a systematic effect of scalp SW origin on the timing of thalamic activity, even though SWs were reported in previous literature (Massimini et al. 2004; Menicucci et al. 2009; Nir et al. 2011; Ujma et al. 2018) and observed in the current study to be local propagating events with a predominantly frontal origin.

Our findings about the primacy of slow activity on scalp channels are in line with previous findings and theories which state that SWs are primarily of cortical origin and while they may appear simultaneously even in subcortical structures, they only do this with a time lag (Nir et al. 2011). A previous study (Mak-McCully et al. 2017) reported similar findings specifically for the pulvinar thalamus: in both, cortical downstates preceded thalamic downstates. We extend this finding to novel thalamic regions: the bilateral anterior thalamus, the mediodorsal nucleus and the ventral anterior nucleus. In all of these regions, slow (delta-alpha) activity tended to follow similar activity on scalp channels by 10-100 milliseconds. Thus, our findings suggest that thalamic downstates are generally the result of propagating cortical neuronal activity patterns with a pattern that appears in all nuclei.

Similarly, our findings about sigma-frequency activity supports previous theories about the role of the thalamus in initiating and maintaining spindle rhythms in the cortex (Fernandez and Lüthi 2020; Steriade 2003), and they are broadly in line with a previous pulvinar study (Mak-McCully et al. 2017) which found that sleep spindles initiate in the thalamus while the downstates are still observed in the cortex. We also found that sigma-frequency activity appeared within thalamic nuclei 50-100 msec earlier than on the scalp, without substantial between-nuclei differences. Our findings suggest that, unlike in case of slow waves, spindle-frequency activity exists in the thalamus before it is observed on the scalp, highlighting the role of the reticular thalamic pacemaker in establishing thalamocortical neuronal firing patterns which contribute to the appearance of the cortical spindles recorded on the scalp (Fuentealba and Steriade 2005; Rovo et al. 2014; Vantomme et al. 2019). This pattern is again not topographically restricted, but appears as a general trend in all nuclei.

Nevertheless, we observed several patterns which were not predicted by previous theory and not reported by earlier studies.

First, while we did find early slow activity on the scalp and early sigma activity in the thalamus, the time lags were modest (~100 msec at the most). This is in contrast to the most relevant previous study (Mak-McCully et al. 2017) who found that pulvinar spindles precede cortical spindles by ~250 msec, and in fact thalamic spindles occur already during the end of cortical downstates. Our results show a much more modest shift between the thalamus and the cortex in the time course of slow and fast frequencies. We note that differences in the investigated thalamic nuclei (pulvinar or the set of nuclei included in our study) are a possible explanation for this discrepancy. Our findings also stand in contrast to another previous study (Tsai et al. 2010), which found that ANT spindles lag scalp spindles. While this nucleus was also covered by our investigation, the Tsai et al study used a smaller sample (N=3), did not anchor sleep spindle activity to scalp-recorded SW downstates and estimated lags indirectly using phase slope indices. These methodological differences may explain the discrepancy, but further studies may also investigate whether sleep spindles not anchored to SWs and thus unaffected by the strongly synchronous neuronal firing patterns observed at this time exhibit a different corticothalamic temporal pattern.

Second, we found that the scalp region of SW origin had little effect on the time course of thalamic activity, in spite of the fact that cortical SWs are well-known propagating activity patterns, generally with a frontal origin (Massimini et al. 2004; Menicucci et al. 2009; Nir et al. 2011; Ujma et al. 2018), a pattern which was empirically replicated in the present sample. If the waxing-waning pattern of synchronous neuronal firing responsible for SWs (Csercsa et al. 2010; Nir et al. 2011; Tononi and Cirelli 2014) were to propagate into the thalamus from a specific cortical area, then it would be expected to reach the thalamus earlier if it originated in this area and later if it had to propagate there from a different cortical region. This is not what we observed here: the time course of SW-locked thalamic activity does not vary much by SW origin, even if we restrict the pool of SWs in each area to those originating from there. We hypothesize that the highly synchronous neuronal firing patterns responsible from SWs do not reach the thalamus through an anatomically circumscribed corticothalamic projection with a fixed cortical origin, but instead simultaneously, immediately start propagating towards both other cortical and thalamic areas wherever they first appear. This hypothesis has some support in previous literature: a study of anesthetized mice (Sheroziya and Timofeev 2014) found that SW-related activity starts simultaneously in the cortex and in several thalamic nuclei, suggesting that large-scale cortical activity does not need to occur before the neuronal firing patterns characteristic for SWs also appear in the thalamus.

Third, we also found that thalamic nuclei all exhibit similar local field potentials during scalp SWs, without substantial differences in the frequency or time course of observable activity. This is significant because thalamic nuclei differ greatly in their characteristic innervation patterns (Cappe et al. 2009; Saalmann 2014). However, it is likely that cortical down- and upstates do not reach the thalamus through these main (often subcortical) afferents which may be far removed from the cortex. If this was the case, we would expect a substantially greater time lag between cortical and thalamic slow activity and much more heterogeneity between the nuclei due to the different degree of indirectness of their afferents from the cortex. The presence of nearly simultaneous slow activity in the cortex and all recorded thalamic nuclei suggests that this activity from the cortex propagates into the thalamus in a relatively direct and possibly nucleus-aspecific manner, circumventing the main afferents. Thus, the great degree of functional differentiation between the thalamic nuclei (Cappe et al. 2009) may be short-circuited in NREM sleep, with neurons in at least most of the thalamus tuned to a rhythm dictated by external pacemakers. NREM sleep oscillations propagating through a relatively direct corticothalamic communication pathway would also explain the generally high synchrony between thalamic and cortical rhythms (Tsai et al. 2010) and the high degree of co-occurrence between cortical and thalamic spindles (Bastuji et al. 2020) observed across several thalamic nuclei.

Our work suffers from a number of limitations. First, our methodology relied on a precise estimation of the temporal offset between scalp SWs and thalamic activity and required a clean, stereotypical thalamic waveform around the SW peak. Our study was not designed to discover millisecond-scale differences between the time course of time-frequency activity across different sources and scalp region or thalamic nucleus differences of this magnitude may have remained hidden. However, differences of at least a few dozen milliseconds (that is, in the range of what was found in scalp vs. thalamus comparisons) would have been readily discoverable, but were not seen. Second, for the same reason our results must be interpreted with caution where a high level of precision was not guaranteed due to insufficient data or the lack of coupling: specifically, in case of the VA nuclei where only data from a maximum of two patients was available, in case of original SWs where the number of SWs in some scalp channels and in case of the gamma band where activity was not systematically coupled to SWs. Third, a general caveat in all human invasive EEG studies is that it is unknown to what degree our findings were biased by the underlying epileptic pathology of our research participants.

In sum, our work provides evidence for a widespread, stereotypical neuronal activity pattern in the human thalamus during scalp SWs, with little heterogeneity across thalamic nuclei or scalp origins. While slow activity occurs first on the scalp and fast activity first in the thalamus, time lags are modest and the time-frequency composition of local thalamic field potentials is similar to what is observable on the scalp. We hypothesize that slow rhythms in the thalamus are strongly affected by propagating cortical activity through nonspecific corticothalamic projections, but we also find evidence for a reverse pattern in case of spindle-frequency activity.

## Acknowledgements

The project was supported by the National Research, Development and Innovation Office of Hungary (NKFI_FK_128100, K_128117 and NKFIH-1157-8/2019-DT), as well as by the Higher Education Institutional Excellence Program of the Ministry of Human Capacities in Hungary, within the framework of the Neurology thematic program of the Semmelweis University.

## Data availability

Supplementary data, raw data (frequency bin and time point resolution statistics) and the MATLAB codes used for analyses are available on Zenodo at the following address: 10.5281/zenodo.5560997

Further data is available upon reasonable request to the Corresponding author.

## Notes

### Competing Interest Statement

The authors have declared no competing interest.

