## Supplementary figure S1 for "Thalamic activity during scalp slow waves in humans"

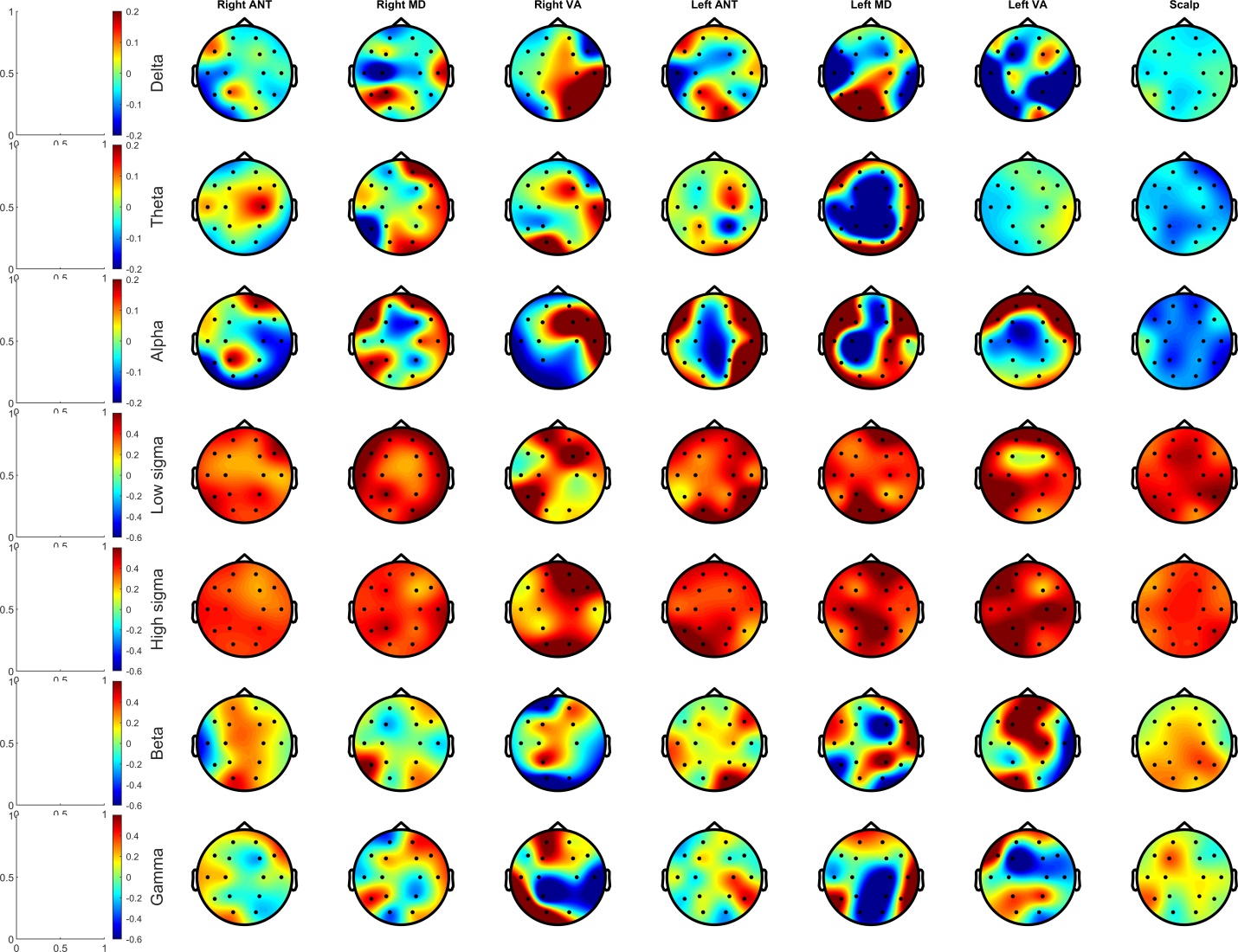

Supplementary figure S1. Temporal lag of activity in each frequency band relative to SW negative peaks as a function of thalamic nucleus and scalp region. Only SWs originating from each channel were considered for this analysis. Data from the scalp itself is shown for reference.
